# A di-acidic motif targets cytoplasmic proteins for unconventional protein secretion

**DOI:** 10.1101/122028

**Authors:** David Cruz-Garcia, Nathalie Brouwers, Juan M. Duran, Amy J. Curwin, Vivek Malhotra

## Abstract

We previously reported that Acb1, a cytoplasmic protein in *Saccharomyces cerevisiae* that cannot enter the endoplasmic reticulum (ER), was secreted upon nutrient starvation by a Vps4 independent, but ESCRT-I, -II and -III and Grh1 dependent pathway (Curwin et al., 2016). Here, we report that the same conditions result in secretion of another signal sequence lacking protein, superoxide dismutase 1 (SOD1). Similar to Acb1, SOD1 export requires Grh1 and a subset of ESCRT components. Importantly, our analysis reveals the existence of a conserved di-acidic motif (Asp-Glu) in SOD1 and Acb1 that is required for their respective export. This sequence is different from the di-acidic motif (Asp-X-Glu) necessary for export of transmembrane proteins from the ER. We propose that the Asp-Glu sequence acts as a targeting motif for the entry of SOD1 and Acb1, and likely many other proteins, upon nutrient starvation into a common albeit ER-Golgi independent pathway of secretion.

## Introduction

A large number of proteins that lack a signal sequence for entering the conventional endoplasmic reticulum (ER)-Golgi complex pathway of secretion are exported by eukaryotic cells (Kinseth et al., 2007; Rabouille et al., 2012; Malhotra, 2013). The challenge is to reveal the mechanism and physiological significance of this unconventional mode of secretion.

In search of a function for GRASP proteins, we discovered that deletion of the single GRASP gene in *Dictyostelium discoideum* blocked starvation-specific secretion of signal sequence lacking acyl CoA binding protein AcbA (Kinseth et al., 2007). This rekindled interest in this unconventional mode of secretion since its first description that signal sequence lacking interleukin (IL)-1ß was secreted by activated macrophages (Auron et al., 1984; Rubartelli et al., 1990). Subsequent analysis revealed that the function of GRASP orthologs (Grh1 in budding yeast) in the secretion of AcbA ortholog (Acb1 in yeast) was conserved in eukaryotes (Duran et al., 2010; Manjithaya et al., 2010). GRASP proteins are also shown to have a role in secretion of signal sequence lacking IL-1ß (Dupont et al., 2011; Zhang et al., 2015), and insulin-degrading enzyme (Son et al., 2016). Altogether, these data lend confidence in the involvement of GRASP proteins in unconventional secretion. The Grh1 dependent export of Acb1 in yeast involves ESCRT-I, -II, and -III. Surprisingly, however, ESCRT mediated secretion of Acb1 is independent of Vps4, which strongly suggest that its secretion is without incorporation into intraluminal vesicles at multivesicular bodies (Curwin et al., 2016). The secretion of Acb1 is therefore independent of exosome-mediated release of cytoplasmic contents. Autophagy related proteins are also suggested to play a role in unconventional protein secretion, but it is unclear whether the dependence on Atg genes is direct or if they are required to keep cells in a response-competent status under nutrient starvation or stress conditions that result in the release of proteins such as Acb1 and IL-1ß (Duran et al., 2010; Manjithaya et al., 2010; Dupont et al., 2011; Zhang et al., 2015).

Conditions that trigger secretion of Acb1 result in the biogenesis of a new Grh1 containing compartment called CUPS that appears in two sequential forms (Bruns et al., 2011; Curwin et al., 2016). Grh1 is first observed in a cluster of vesicles that are produced independently of COPII-and COPI-mediated export of membranes from the ER and the Golgi, respectively. Membrane export from late Golgi compartment, however, is necessary for production of these vesicular elements (Bruns et al., 2011; Cruz-Garcia et al., 2014). Cluster of Grh1 containing vesicles are then found surrounded by curved saccules (Curwin et al., 2016). The stability of this form of CUPS is dependent on the activity of the phosphatidylinositol 3-phosphate kinase Vps34 (Cruz-Garcia et al., 2014). For the sake of simplicity we have decided to call the first stage of CUPS: I-CUPS for immature CUPS (cluster of Grh1 containing vesicles), and the second as M (mature)-CUPS (Grh1 containing vesicles surrounded by curved saccules). Acb1 is associated with M-CUPS indicating their involvement in events leading to Acb1 secretion (Curwin et al., 2016).

The next obvious challenge, in order to understand the mechanism of unconventional protein secretion, is to identify whether there is a motif – a sequence – used by proteins to enter this pathway. An easy means to address this concern is to ask how many other proteins are secreted by the same pathway and then to test whether such proteins contain a common sequence, which when mutated affects their secretion. We now present data that exactly addresses this issue. We report that signal sequence lacking cytoplasmic superoxide dismutase 1 (SOD1) is exported by the pathway of Acb1 secretion. In addition, we reveal that SOD1 and Acb1 contain a di-acidic motif that is necessary for their release from cells. The description of our findings follows.

## Results and Discussion

### SOD1 and Acb1 follow the same ER-Golgi independent secretory pathway

We have previously shown that Acb1 is secreted by yeast incubated in 2% potassium acetate for 2 to 3 hr. Extraction of the yeast cell wall is used to quantitate the secreted pool of Acb1. Lack of cofilin-1, an actin filament interacting protein, in the extracted material serves as a control for release of cytoplasmic content as a result of cell lysis during experimental manipulations (Curwin et al., 2016). In addition, presence of the cell wall protein Bgl2 is used as a control to monitor extraction efficiency in this assay (Curwin et al., 2016).

The cytosolic copper-zinc superoxide dismutase SOD1, which lacks a signal sequence, is released by a number of different mammalian cells (Mondola et al., 1996, 1998; Cimini et al., 2002). Although the mechanism of SOD1 export by cells is unclear, it has been reported that rat pituitary GH3 cells secrete SOD1 in response to depolarization through SNARE dependent exocytosis, which allowed the authors to suggest the involvement of large dense-core vesicles containing SOD1 (Santillo et al., 2007). SOD1 has also been found associated with exosomal fraction obtained from tumor cell lines in high-throughput studies (ExoCarta database; www.exocarta.org/). Furthermore, immunochemical analysis has revealed that a fraction of extracellular human SOD1 is associated with exosomes secreted by mouse astrocytes and mouse motor neurons overexpressing wild-type human SOD1 (Gomes et al., 2007; Basso et al., 2013; Grad et al., 2014). Therefore, SOD1 is released by cells and it might be associated with a membrane bounded compartment, but the conditions that trigger its release and the underlying mechanism are largely unknown.

Budding yeast also express a signal sequence lacking SOD1 of 16 kDa. Therefore, we tested whether SOD1 was secreted when cells are grown in nutrient-rich conditions or incubated in potassium acetate. Yeast were grown to mid-logarithmic phase in synthetic complete medium or cultured in 2% potassium acetate for 2.5 hr. After collecting the cells, the cell wall was extracted as described previously and western blotted with specific antibodies to detect Acb1, SOD1, cofilin-1 and Bgl2 (Curwin et al., 2016). We detected a moderate amount of SOD1 in the cell wall of yeast grown in nutrient-rich medium (Figure 1A). However, cells incubated in potassium acetate revealed a 9-fold increase in cell wall extractable SOD1 compared to control cells (Figure 1A-B). This level of starvation-induced secretion of SOD1 is similar to that reported previously, and shown here, for Acb1 (Curwin et al., 2016; Figure 1A). Cofilin-1 was not detected in the cell wall extracts obtained from starved cells or cells cultured in nutrient-rich medium (Figure 1A). This data indicates that, like Acb1, SOD1 secretion by yeast cells is stimulated by starvation in potassium acetate.

**Figure 1.**
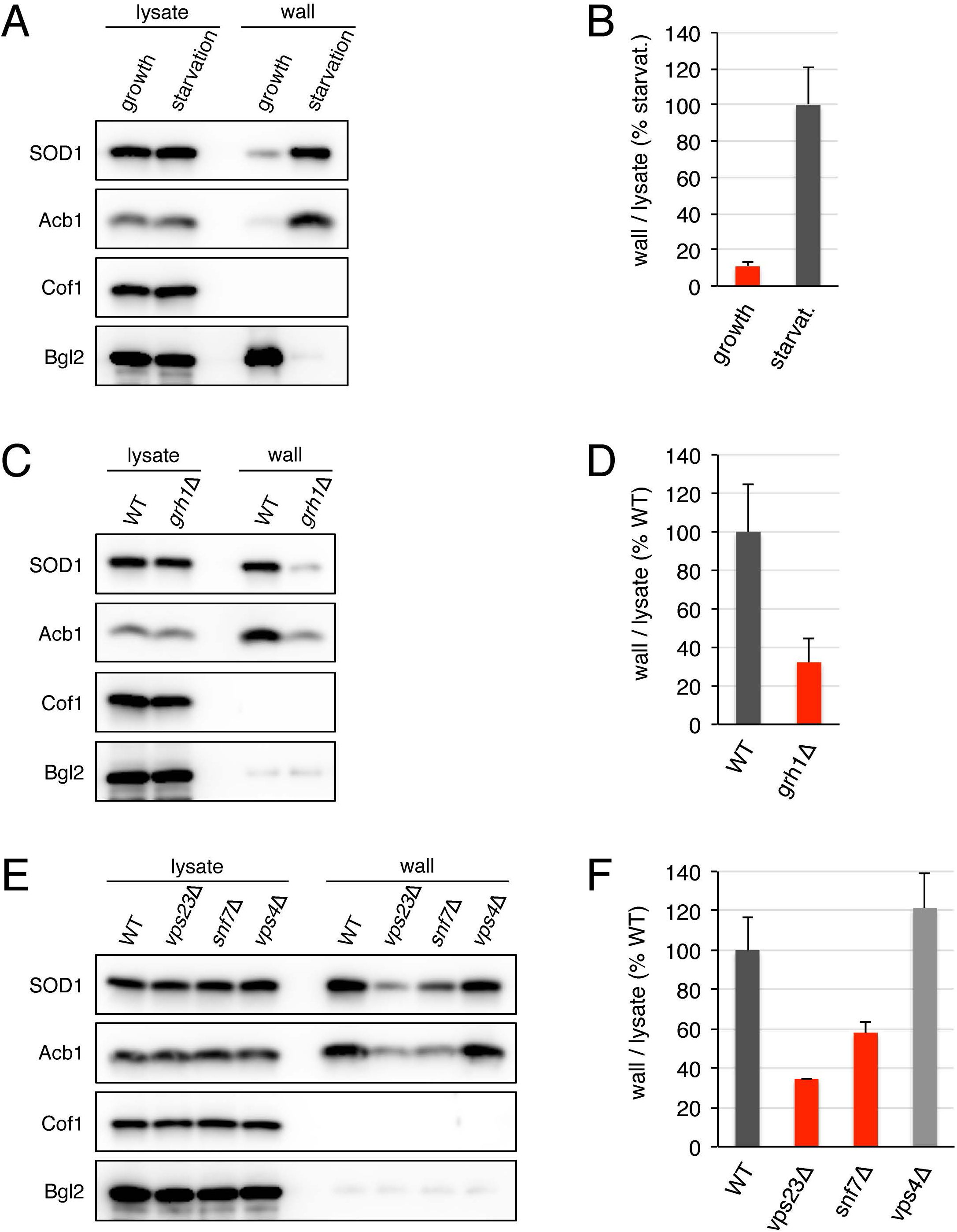
SOD1 and Acb1 are secreted by the same pathway in response to nutrient starvation. (A-B) Secretion of SOD1 is induced by nutrient starvation. Wild-type cells were grown in nutrient-rich medium to mid-logarithmic phase (growth). An aliquot of this culture was washed twice, and incubated in 2% potassium acetate for 2.5 hr (starvation). Cell wall proteins were extracted from equal numbers of growing and starved cells followed by precipitation with TCA. Lysates and cell wall-extracted proteins (wall) were analyzed by western blot. SOD1 levels were quantified, the ratio of wall/lysate SOD1 was determined, and represented as percentage of the starved wild-type culture. (C-D) Grh1 is required for SOD1 secretion. The ratio of wall/lysate SOD1 in wild-type and *grh1Δ* cells incubated in 2% potassium acetate for 2.5 hr was determined as in (A-B). (E-F) SOD1 secretion requires a subset of ESCRT proteins. The ratio of wall/lysate SOD1 in wild-type, *vps23Δ, snp7Δ, and vps4Δ* cells incubated in 2% potassium acetate for 2.5 hr was determined as in (A-B). Results are shown as the mean±s.e.m. of three experiments.

Secretion of Acb1 depends on Grh1 and a subset of ESCRT components (Curwin et al., 2016) and we tested their involvement in potassium acetate-induced SOD1 export. In GRH1 deleted cells, the level of SOD1 secreted to the cell wall after 2.5 hr of starvation in potassium acetate decreased by 68% compared to wild-type strain (Figure 1C-D). Similarly, yeast strains deleted of either VPS23 (ESCRT-I) or SNF7 (ESCRT-III) showed a 66% and 52% reduction, respectively, in the levels of cell wall-associated SOD1 as compared with the wild-type control (Figure 1E-F). However, as reported previously for the lack of requirement of Vps4 in Acb1 secretion, deletion of VPS4 did not have a significant effect on SOD1 export and if anything loss of Vps4 slightly increased the pool of secreted SOD1 (Figure 1E-F). So the quantitative effects of the deletions of GRH1 and the selected ESCRT genes on starvation-induced SOD1 secretion are similar to those previously reported for Acb1 release and reconfirmed here (Curwin et al., 2016; Figure 1C and 1E). These findings thus reveal that a new protein, SOD1, requires the same set of components than Acb1 for its export upon starvation.

### Identification of a di-acidic motif essential for starvation-induced SOD1 secretion

Eukaryotes are known to express two highly related classes of copper-zinc superoxide dismutase enzymes that play widespread roles in oxidative stress resistance and signalling. These two classes include the intracellular, predominantly cytosolic, SOD1 and the extracellular SOD (EC-SOD), that contains a signal sequence for entering ER, thereby allowing its secretion via the conventional secretory pathway (reviewed in Zelko et al., 2002). Sequence and structural analysis have revealed that EC-SOD and SOD1 diverged early in evolution (Zelko et al., 2002). In fact, the structural core of the ancestral SOD1 is highly conserved in EC-SOD and consists of a β-barrel motif with two large loops: the so-called electrostatic loop and the zinc-binding loop (Figure 2A-B; Zelko et al., 2002). We noticed that, within this structural core, EC-SOD and SOD1 share high sequence homology with the exception of a short stretch of 8 amino acids that is contained in SOD1 but not in EC-SOD (Figure 2B-C). This stretch (amino acids: 73-GGPKDEER-80 in the sequence of human SOD1) is located in the zinc-binding loop and flanked by the conserved histidines H72 and H81, which coordinate the zinc cation (Figure 2B-C). Taking into account that this short stretch is conserved in the SOD1 sequences from yeast to human and that it has been lost in the sequences of vertebrate EC-SOD, we hypothesized that it might be important for the unconventional secretion of SOD1.

**Figure 2.**
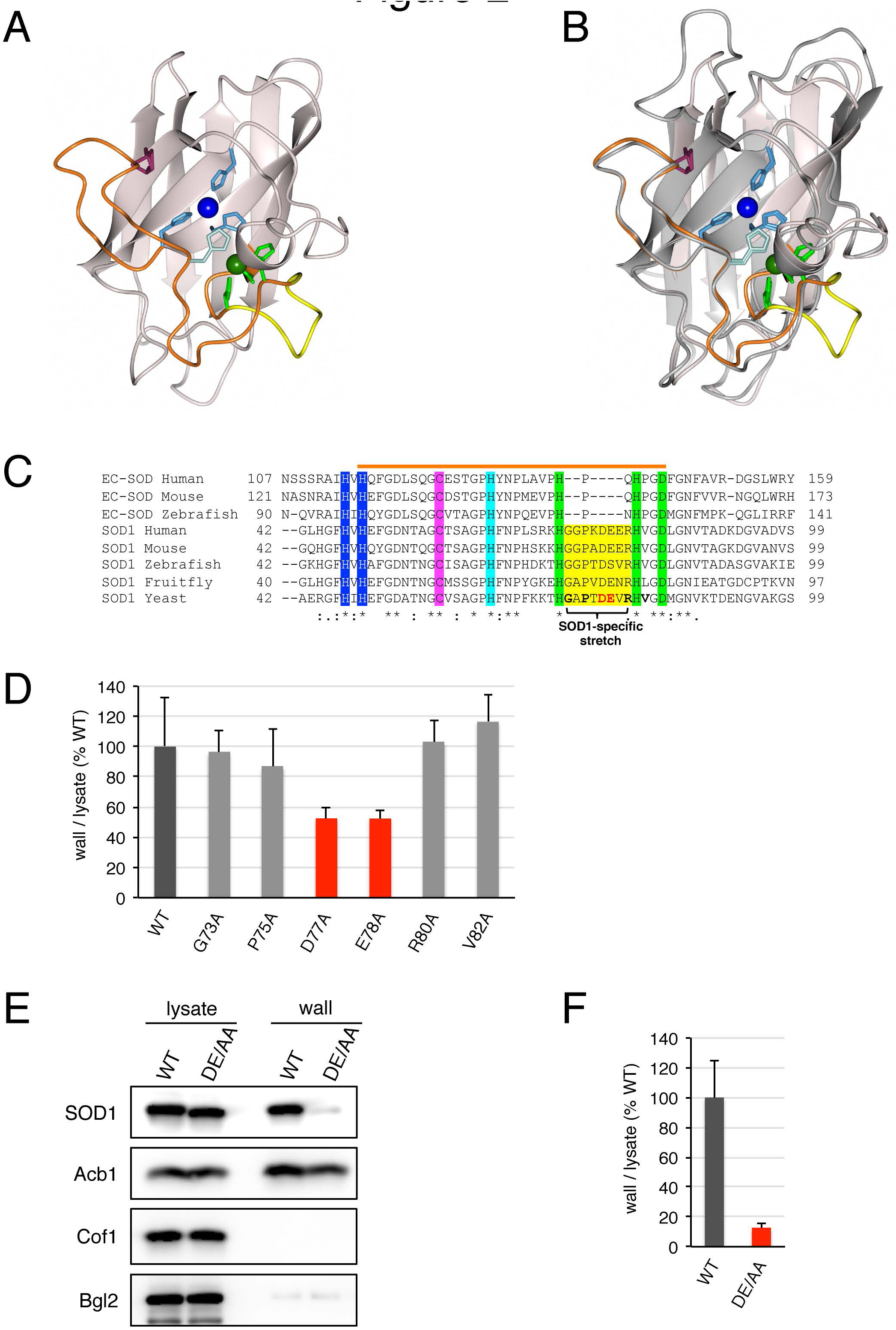
Identification of a di-acidic sequence in SOD1 for its export. (A) 3D structure of human SOD1 monomer (PDB: 2C9U). The β-barrel is shown in pink; copper is in dark blue; zinc is in dark green; the intramolecular disulfide bond is shown in purple; zinc-binding loop is in orange; in light blue are the side chains of H47, H49 and H121; the side chains of H72, H81 and D84 are in light green; H64 is in turquoise; and the SOD1-specific amino acid stretch is shown in yellow. (B) Superimposition of the 3D structures of human SOD1 monomer and human EC-SOD monomer (PDB: 2JLP). Structural features of human SOD1 are shown as in (A). Human EC-SOD monomer is shown in grey. Side-chains coordinating copper and zinc in human EC-SOD are colour-coded as in (A). (C) Alignment of partial amino acid sequences of human, mouse, and zebrafish EC-SOD, and human, mouse, zebrafish, fruitfly, and budding yeast SOD1. The zinc-binding loop is overlined with an orange bar; copper-binding histidines are in blue; residues coordinating zinc are in green; the histidine bridging copper and zinc is in turquoise; one of the cysteines forming the intramolecular disulfide bond is in purple; the SOD1-specific stretch is yellow; the residues in yeast SOD1 mutated to alanine in this study are shown in bold; the di-acidic DE77/78 motif in yeast SOD1 is in bold red letters. (D) Effect of single alanine substitutions in the SOD1-specific stretch on SOD1 secretion. Cells expressing either wild-type SOD1 or single alanine substitutions in positions G73, P75, D77, E78, R80, or V82 of SOD1 were grown to mid-logarithmic phase, washed twice, and incubated in 2% potassium acetate for 2.5 hr. Cell wall proteins were extracted from equal numbers of cells followed by precipitation with TCA. Lysates and cell wall-extracted proteins were analyzed by western blot. The ratio of wall/lysate SOD1 was determined and represented as percentage of the strain expressing wild-type SOD1. (E) The DE77/78 di-acidic motif is essential for SOD1 secretion. Representative cell wall extractions monitored by western blot of either wild-type or DE77/78AA SOD1 expressing cells incubated for 2.5 hr in 2% potassium acetate. (F) SOD1 levels were quantified, the ratio of wall/lysate SOD1 was determined, and represented as percentage of the starved wild-type culture. Results are shown as the mean±s.e.m. of at least three experiments.

We tested this hypothesis using nutrient starvation-induced SOD1 secretion by yeast cells. Budding yeast only express SOD1 and lacks EC-SOD. We generated yeast strains where the five residues within this short stretch that are conserved between yeast and human SOD1 (G73, P75, D77, E78, and R80) were individually replaced with alanine in the genomic locus of SOD1 (Figure 2C). In addition, we substituted V82 to alanine, which is very close to this stretch of residues and is conserved in the sequences of SOD1 from yeast to human (Figure 2C). As shown in Figure 2D, substituting G73, P75, R80, or V82 with alanine did not affect starvation-induced secretion of SOD1. However, replacement of either D77 or E78 to alanine reduced the export of SOD1 in response to nutrient starvation by 48% with respect to the wild-type SOD1 strain (Figure 2D). The double substitution DE77/78AA strongly decreased the secreted pool of SOD1 by 88% as compared to the wild-type cells (Figure 2E-F). Importantly, the levels of Acb1 in the cell wall fraction were not affected by the DE77/78AA mutation in SOD1 (Figure 2E) indicating that the pathway of unconventional secretion is fully induced in response to nutrient starvation in the strain expressing the SOD1-DE77/78AA mutant. Moreover, the intracellular levels of SOD1-DE77/78AA mutant were similar to the wild-type SOD1 suggesting that the stability of SOD1 was unaffected by these mutations (Figure 2E). This therefore reveals, for the first time, a sequence in SOD1 that is required for its export from cells upon nutrient starvation.

Next, we investigated whether the DE77/78AA mutation alters the intracellular distribution of SOD1. Examination by fluorescence microscopy revealed that SOD1-DE77/78AA mutant tagged with GFP displays the same diffuse cytoplasmic distribution than the wild-type SOD1 (Figure 2-figure supplement 1), suggesting that the DE77/78AA substitution does not affect the intracellular location of SOD1.

### Acb1 also requires a di-acidic motif for its export

We next asked whether a similar di-acidic motif was required for starvation-induced export of Acb1. Budding yeast Acb1 contains a single DE motif at positions 23-24, which is largely conserved from yeast to human (Figure 3A). This motif is located just at the beginning of the second α-helix in the 3D structure of human Acb1 ortholog called ACBP (Figure 3A). We generated a strain where DE23/24 residues were substituted by alanine in the genomic locus of ACB1. Using the cell wall extraction procedure described above, we found that the levels of Acb1 secreted to the cell wall after 2.5 hr of starvation were 42% lower in the Acb1-DE23/24AA strain compared to the control strain expressing wild-type Acb1 (Figure 3B-C). Therefore, a di-acidic motif is also required for efficient secretion of Acb1 in response to nutrient starvation. The starvation-induced secretion of wild-type SOD1 was unaffected by the DE23/24AA mutation in Acb1 (Figure 3B).

**Figure 3.**
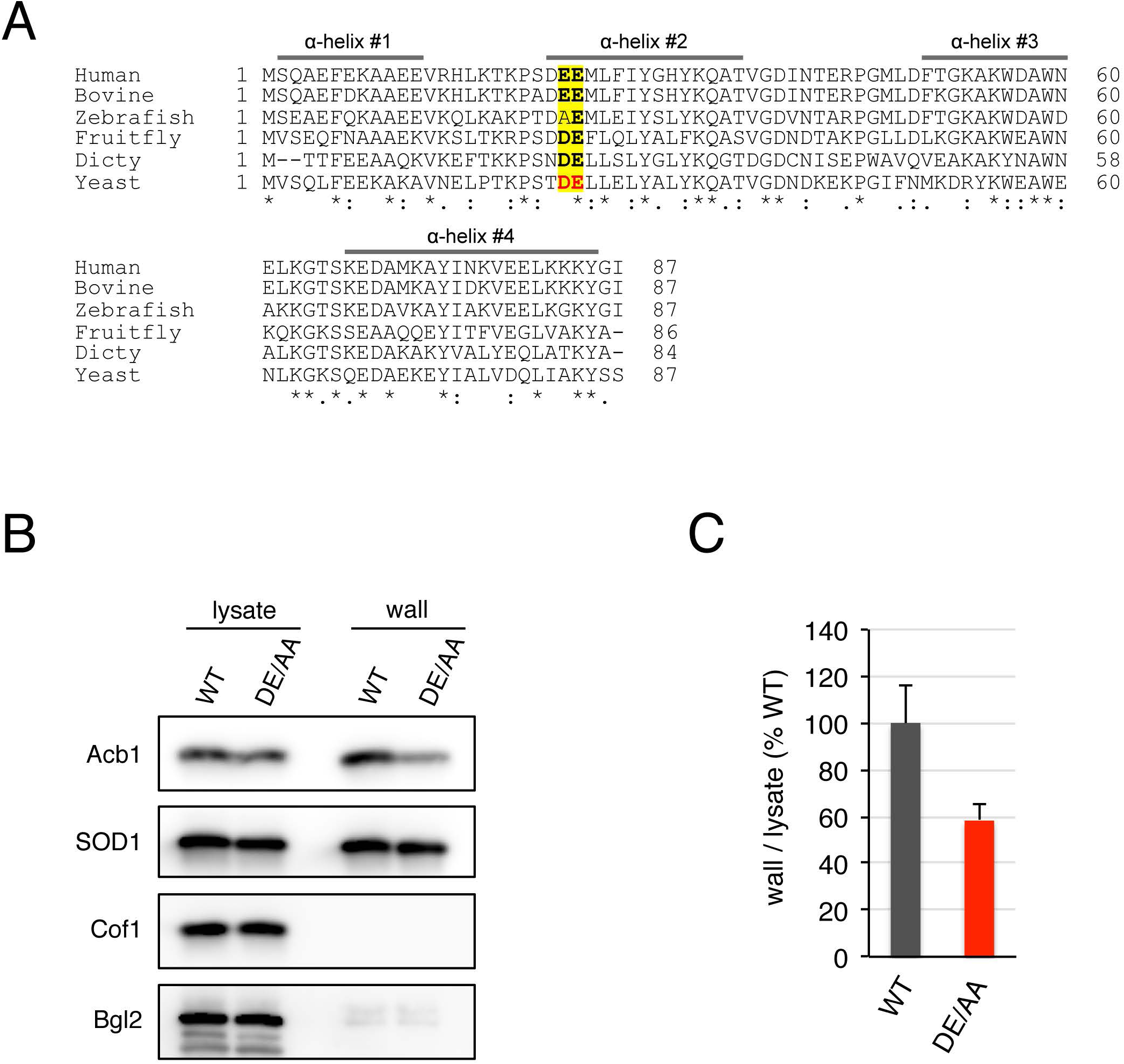
The di-acidic motif DE23/24 of Acb1 is required for its secretion (A) Amino acid sequence alignment of human, bovine, zebrafish, fruitfly, *D. discoideum*, and budding yeast Acb1 orthologs. The four α-helices of human ACBP are indicated by bars; the di-acidic DE23/24 motif in yeast Acb1 is in bold red letters; the position of this di-acidic motif in Acb1 orthologs is indicated in yellow; the conserved residues within the di-acidic motif are in bold letters. (B) The DE23/24 sequence is required for efficient Acb1 secretion. Representative cell wall extractions monitored by western blot of either wild-type or DE23/24AA Acb1 expressing cells incubated for 2.5 hr in 2% potassium acetate. (C) Acb1 levels were quantified, the ratio of wall/lysate Acb1 was determined, and represented as percentage of the starved wild-type culture. Results are shown as the mean±s.e.m. of four experiments.

Based on the data presented here, we suggest that the di-acidic motifs in SOD1 and Acb1 act as targeting sequences for their export upon nutrient starvation. To our knowledge this is the first description of a sequence required for ER independent export and shared by proteins that follow the same pathway.

Interestingly, the negatively charged side-chains of the residues of the di-acidic motif face the solvent in the 3D structure of both human SOD1 and human ACBP (Figure 3-figure supplement 1A-B). It is therefore likely that this motif binds specific proteins that promote their capture for export by the ER independent pathway of secretion. It is important to note that the di-acidic motif in Acb1 and SOD1 (D-E) is different from the di-acidic motif (D-X-E) that captures and exports transmembrane proteins by Sec24 of COPII-coated vesicles at the ER (Barlowe, 2003). Moreover, COPII coats are not involved in the export of unconventionally secreted cargoes like Acb1 (Duran et al., 2010; Cruz-Garcia et al., 2014). The finding of this di-acidic motif for unconventional secretion is clearly an important milestone, but we expect many layers of events that select proteins for this mode of secretion. This is for the following reasons. 1, Only a small fraction of Acb1 and SOD1 are secreted in a nutrient starvation dependent manner and 2, many other cytoplasmic proteins contain this sequence so why are they not secreted along with Acb1 and SOD1. For example, cofilin-1 contains a di-acidic sequence (D-E) at residues 10-11, but is not secreted under conditions that release Acb1 and SOD1. How a di-acidic sequence is exploited by the cells to select only a small fraction of the total cellular mass of the proteins for unconventional secretion is the obvious next challenge and identification of the binding partners of the di-acidic motif could potentially help resolve this issue.

## Materials and Methods

### Media, yeast strains and plasmids

Yeast cells were grown in synthetic complete (SC) media (0.67% yeast nitrogen base without amino acids, 2% glucose supplemented with amino acid drop-out mix; SIGMA-Aldrich, St. Louis, MO, USA). All *Saccharomyces cerevisiae* strains used in this study are listed in Table 1. For deletion of GRH1, the open reading frame (ORF) was replaced with a KanMX4 cassette using PCR-based targeted homologous recombination as previously described (Cruz-Garcia et al., 2014). To generate SOD1 mutant strains, the yeast SOD1 ORF was first PCR amplified from genomic DNA and cloned into the plasmid p416-ADH using BamHI and XhoI sites. From this plasmid, the SOD1 ORF and CYC1 terminator were PCR amplified and introduced into the pFA6a-His3MX6 plasmid using BamHI and XmaI sites. Mutations in the SOD1 ORF were introduced using Gibson assembly (Gibson et al., 2009). The C-terminally GFP-tagged SOD1 wild-type or DE77/78AA mutant constructs were generated by replacing the SOD1 stop codon and CYC1 terminator by a fragment encompassing a linker-GFP-ADH1 terminator. For this the SOD1 ORF was amplified from the respective construct described above and the linker-GFP-ADH1 terminator fragment was PCR amplified from pYM44 (Janke et al., 2004). Both fragments were introduced in pFA6a-His3MX6 by Gibson assembly to obtain the GFP-tagged SOD1 wild-type or DE77/78AA mutant constructs. To generate Acb1 mutant strains, we introduced the His3MX6 cassette into pRS416-PrACB1-Acb1-3xFlag (Duran et al., 2010) by Gibson assembly at the BglII site located in the ACB1 promoter region. Mutations in the ACB1 ORF were introduced by Gibson assembly. All mutant constructs were confirmed by nucleotide sequence analysis covering the coding regions in the construct. Strains expressing mutated SOD1, SOD1-GFP or Acb1 and the corresponding wild-type controls were then generated by PCR-based targeted homologous recombination using the respective constructs. Single colonies were selected and screened for the presence of the respective mutation by nucleotide sequence analysis. SnapGene software (GSL Biotech, Chicago, IL, USA) was used for molecular cloning design.

### Assay for unconventional protein secretion

Yeast cells were inoculated at 0.004–0.008 OD_600_/mL in SC medium and grown overnight at 25°C. The following day, when cells had reached OD_600_/mL of 0.4–0.7, equal numbers of cells (15 OD_600_ units) were harvested, washed twice in sterile water, resuspended in 1.5 mL of 2% potassium acetate and incubated for 2.5 hr. Concomitant to this, growing cells were diluted in SC medium, continued growing in logarithmic phase and 15 OD_600_ units were harvested as before. The cell wall extraction and the TCA precipitation of the extracted material were performed exactly as described earlier (Curwin et al., 2016). For immunoblotting, proteins (10 μL each of lysate or wall fractions) were separated in a 16.5% Tris-tricine peptide gel (Bio-Rad, Hercules, CA, USA) before transfer to nitrocellulose membrane. Rabbit anti-yeast SOD1 antibody was a gift from Dr. Thomas O’Halloran (Northwestern University, IL, USA). Rabbit anti-Cof1 antibody was a generous gift from Dr. John Cooper (Washington University in St. Louis, MO, USA). Rabbit anti-Bgl2 was a gift from Dr. Randy Schekman (UC Berkeley, CA, USA). Rabbit anti-Acb1 antibody has been described elsewhere (Curwin et al., 2016).

### Epifluorescence microscopy

After incubation in the appropriate medium cells were harvested and examined by epifluorescence microscopy using a DMI6000 B microscope (Leica microsystems, Wetzlar, Germany) as described earlier (Curwin et al., 2016). Images were acquired using LAS AF software (Leica microsystems) and processing was performed with ImageJ 1.45r software.

### Bioinformatic analysis

Amino acid sequences were downloaded from UniProt (UniProt Consortium, 2014) and aligned using Clustal Omega (Sievers et al., 2011). All structural figures were created using the CCP4MG program (McNicholas et al., 2011).

## Acknowledgements

We thank members of the Malhotra Lab for valuable discussions. We thank Drs. Thomas O’Halloran (Northwestern University, IL, USA), John Cooper (Washington University, MO, USA), and Randy Schekman (UC Berkeley, CA, USA) for kindly sharing reagents. We acknowledge support from the Spanish Ministry of Economy and Competitiveness through the Programme ‘Centro de Excelencia Severo Ochoa 2013-2017’ (SEV-2012-0208); and support from the CERCA Programme / Generalitat de Catalunya. Vivek Malhotra is an Institució Catalana de Recerca i Estudis Avançats (ICREA) professor at the Center for Genomic Regulation and the work in his laboratory is funded by grants from MINECO’s Plan Nacional (BFU2013-44188-P), Consolider (CSD2009-00016), and European Research Council (268692). The project has received research funding from the European Union. This paper reflects only the author’s views. The Union is not liable for any use that may be made of the information contained therein.

---

**Figure 2-figure supplement 1.**
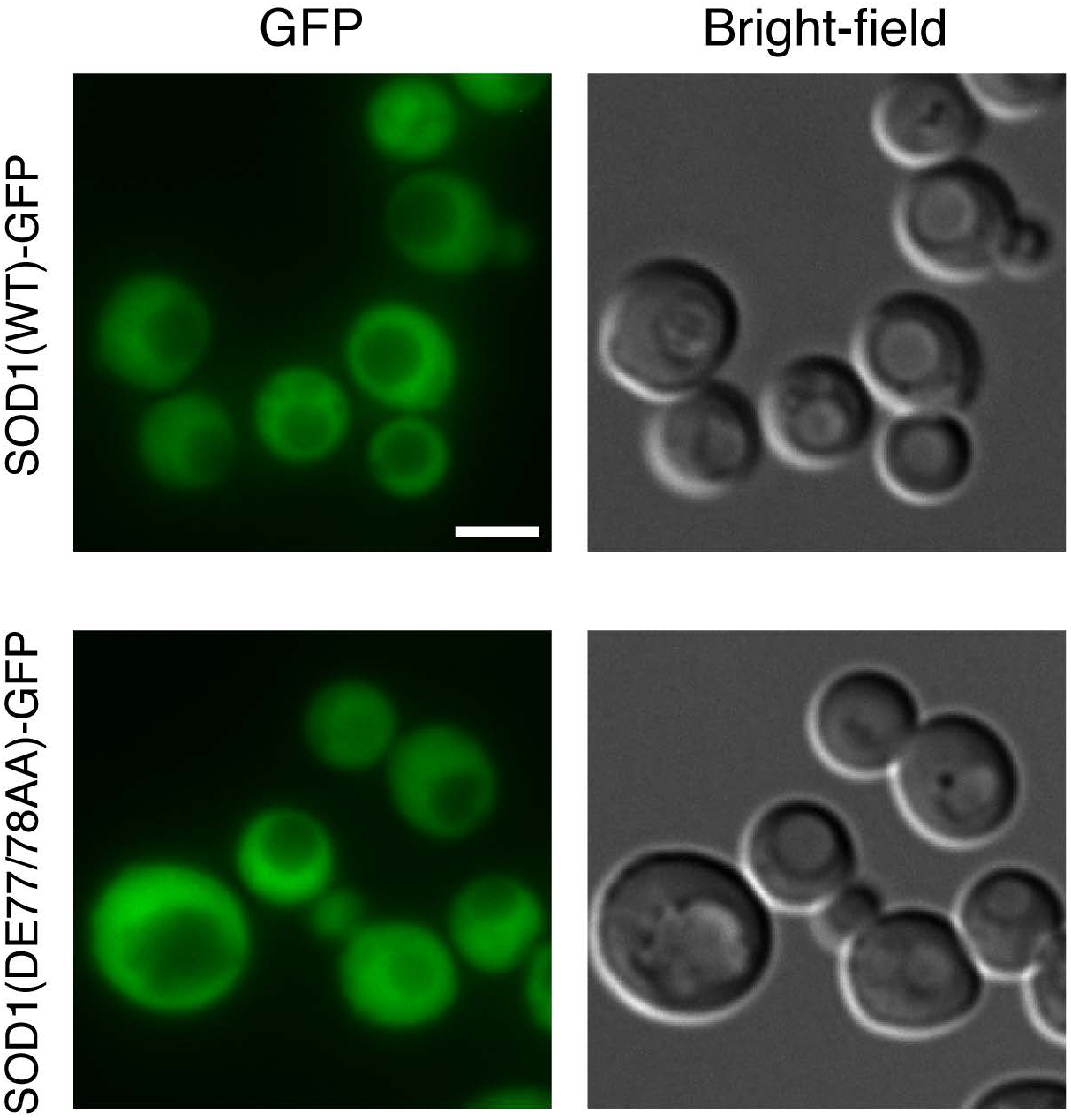
The DE77/78AA mutation does not alter the diffuse cytoplasmic intracellular distribution of SOD1. Cells expressing either C-terminally GFP-tagged wild-type SOD1 or the DE77/78AA mutant of SOD1-GFP were grown to mid-logarithmic phase, washed twice, incubated in 2% potassium acetate for 2 hr, and examined by epifluorescence microscopy. (GFP) Representative micrographs showing the cytoplasmic distribution of wild-type and DE77/78AA SOD1-GFP are presented. Bright-field images are shown for reference purpose. Scale bar = 2 μm.

**Figure 3-figure supplement 1.**
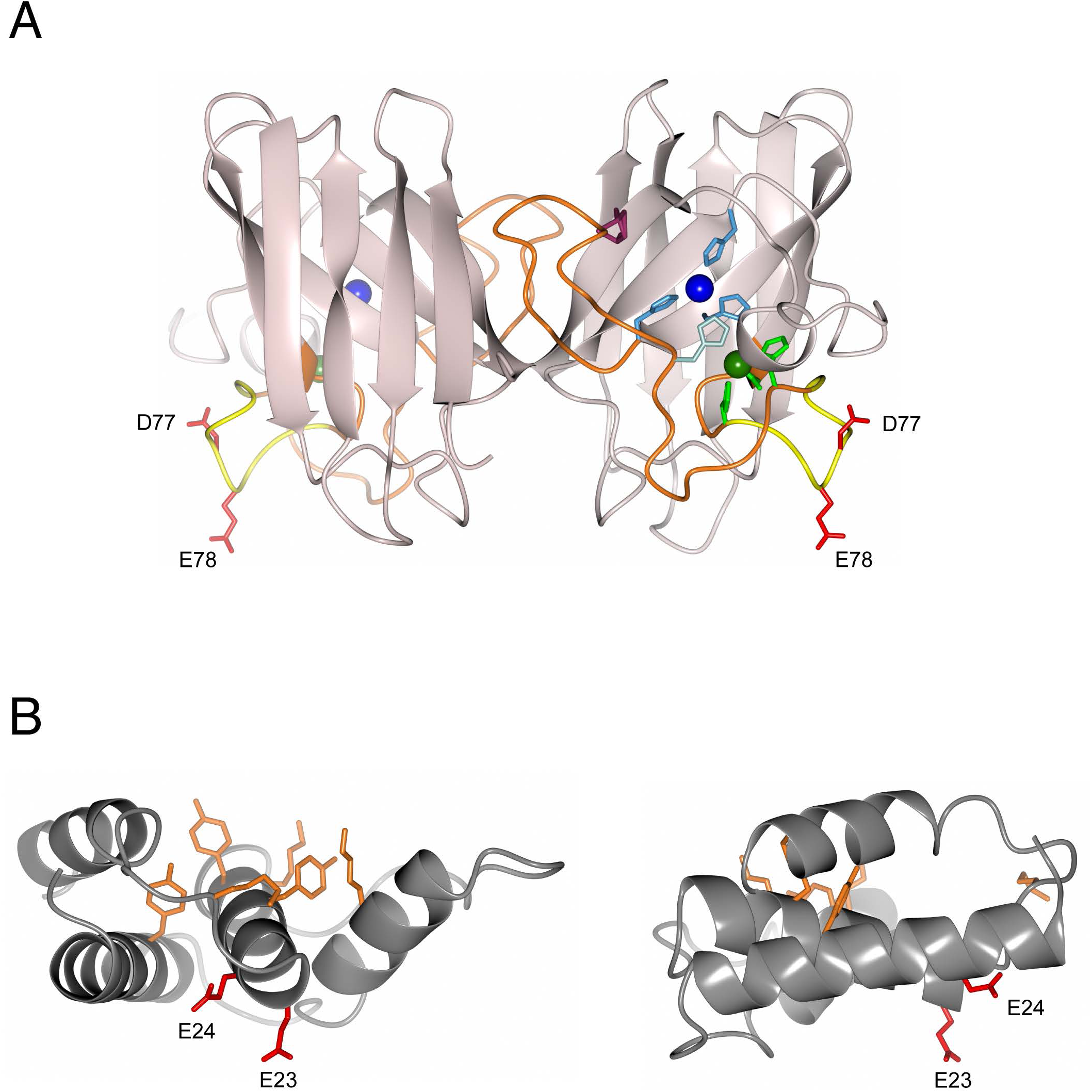
The side-chains of the residues forming the di-acidic motif of SOD1 and ACBP are exposed to the solvent. (A) 3D structure of human SOD1 dimer (PDB: 2C9U). Structural features are shown as in Figure 2A. The side chains of D77 and E78 are in red. (B) Two views of the 3D structure of human ACBP (PDB: 2FJ9). Side chains of residues involved in acyl-CoA-binding (K19, Y29, Y32, K33, K55, Y74) are shown in orange. The side chains of E23 and E24 are in red.

